# Myoglobin Inhibits Breast Cancer Cell Fatty Acid Oxidation and Migration via Heme-dependent Oxidant Production and Not Fatty Acid Binding

**DOI:** 10.1101/2024.04.30.591659

**Authors:** Aaron R. Johnson, Krithika Rao, Bob B. Zhang, Steven Mullet, Eric Goetzman, Stacy Gelhaus, Jesus Tejero, uti Shiva

## Abstract

The monomeric heme protein myoglobin (Mb), traditionally thought to be expressed exclusively in cardiac and skeletal muscle, is now known to be expressed in approximately 40% of breast tumors. While Mb expression is associated with better patient prognosis, the molecular mechanisms by which Mb limits cancer progression are unclear. In muscle, Mb’s predominant function is oxygen storage and delivery, which is dependent on the protein’s heme moiety. However, prior studies demonstrate that the low levels of Mb expressed in cancer cells preclude this function. Recent studies propose a novel fatty acid binding function for Mb via a lysine residue (K46) in the heme pocket. Given that cancer cells can upregulate fatty acid oxidation (FAO) to maintain energy production for cytoskeletal remodeling during cell migration, we tested whether Mb-mediated fatty acid binding modulates FAO to decrease breast cancer cell migration. We demonstrate that the stable expression of human Mb in MDA-MB-231 breast cancer cells decreases cell migration and FAO. Site-directed mutagenesis of Mb to disrupt Mb fatty acid binding did not reverse Mb-mediated attenuation of FAO or cell migration in these cells. In contrast, cells expressing Apo-Mb, in which heme incorporation was disrupted, showed a reversal of Mb-mediated attenuation of FAO and cell migration, suggesting that Mb attenuates FAO and migration via a heme-dependent mechanism rather than through fatty acid binding. To this end, we show that Mb’s heme-dependent oxidant generation propagates dysregulated gene expression of migratory genes, and this is reversed by catalase treatment. Collectively, these data demonstrate that Mb decreases breast cancer cell migration, and this effect is due to heme-mediated oxidant production rather than fatty acid binding. The implication of these results will be discussed in the context of therapeutic strategies to modulate oxidant production and Mb in tumors.

**Graphical Abstract:** Graphical Abstract:
Mb-dependent oxidant generation (but not fatty acid binding) dysregulates mitochondrial respiration and migratory gene expression, leading to decreased cell migration. Created with BioRender.

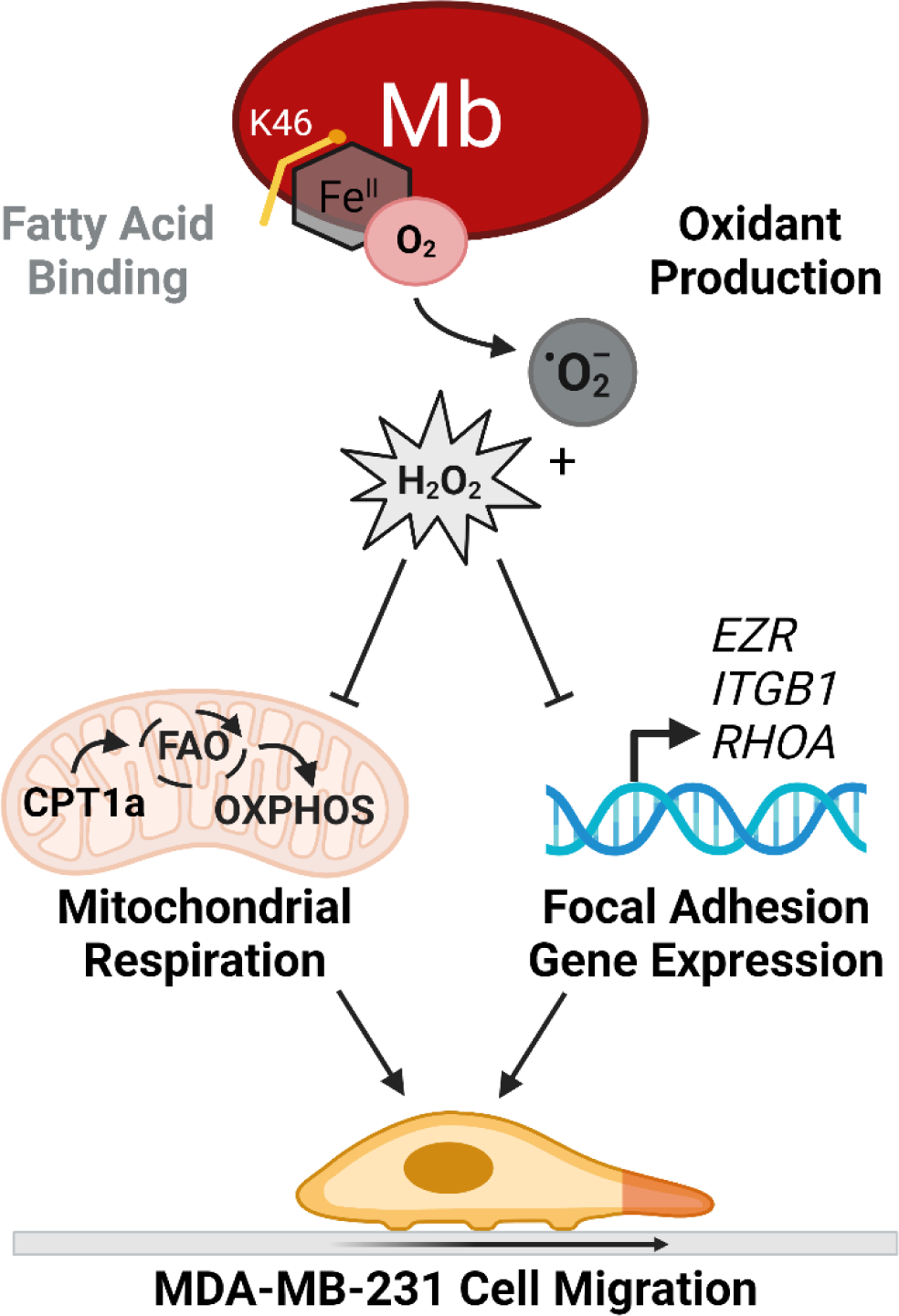

**Highlights:**

- Myoglobin (Mb) expression in MDA-MB-231 breast cancer cells slows migration.
- Mb expression decreases mitochondrial respiration and fatty acid oxidation.
- Mb-dependent fatty acid binding does not regulate cell migration or respiration.
- Mb-dependent oxidant generation decreases mitochondrial metabolism and migration.
- Mb-derived oxidants dysregulate migratory gene expression.

## Introduction

Myoglobin (Mb) is a monomeric heme protein that is constitutively expressed at high concentrations (200-300µM) in skeletal and cardiac muscle, where its canonical role is to store and deliver oxygen to mitochondria to maintain aerobic respiration in hypoxic conditions.^*1*–*6*^ Accumulating studies now demonstrate that Mb can be expressed in several types of human carcinomas, including breast, ovarian, head and neck, prostate, and lung cancers.^*7*–*9*^ In breast cancer patients, Kristiansen *et al.* demonstrated that Mb is ectopically expressed in nearly 40% of all breast tumors.^*10*^ Notably, breast cancer patients with tumor Mb expression show smaller, more differentiated tumors and fewer metastases, which correlate with better survival compared to patients without Mb.^*10*^ However, the mechanisms by which Mb regulates cancer cell metabolism and migration are incompletely understood.

The level of Mb expression within breast tumors (3-10µM) is significantly lower than in myocytes, and others have shown that these low concentrations of Mb are insufficient to mitigate hypoxia in cancer cells.^*11*^ This has prompted the idea that Mb regulates cancer cell function through mechanisms independent of oxygen storage and delivery.^*9*, *11*^ It is well established that in addition to oxygen storage, the heme prosthetic group of Mb functions to regulate nitric oxide (NO•) levels^*12*–*14*^ and catalyze cellular oxidant production and scavenging.^*15*–*17*^ In this regard, we have previously shown that Mb-mediated inhibition of cancer cell proliferation is dependent on its heme-mediated oxidant production,^*15*^ and others have demonstrated that the Mb heme group regulates NO• levels in cancer cells.^*16*, *18*^

A growing body of literature has now demonstrated a novel function for Mb in which the protein binds to several medium- and long-chain fatty acids.^*5*, *19*–*23*^ Computational modeling has identified that the lysine 46 residue (K46) of human Mb stabilizes fatty acids near the heme pocket, and this binding enhances fatty acid oxidation (FAO) in cardiomyocytes.^*24*–*26*^ In MDA-MB-468 and MCF7 breast cancer cells, Armbruster *et al.* showed that endogenous Mb expression increases cytoplasmic solubility of fatty acids while limiting FAO.^*5*^ However, this study did not test whether these effects are directly dependent on Mb-fatty acid-binding. Notably, FAO is upregulated in some breast cancers due to its high yield of ATP and reducing equivalents (e.g., NADH, FADH_2_) per molecule of substrate, which can drive efficient actin cytoskeleton remodeling, promote cell migration, and subsequent metastatic spreading throughout the body.^*27*–*30*^ Despite this recognition, it is unknown whether Mb-dependent fatty acid binding directly regulates FAO and cell migration in breast cancer cell models.

Herein, we tested whether Mb binds fatty acids to regulate fatty acid metabolism and cell migration in breast cancer cells. We find that Mb-dependent fatty acid binding does not affect mitochondrial FAO or cell migration. Instead, our data demonstrate that Mb’s heme-dependent oxidant production inhibits breast cancer cell FAO and migration. The implication of these results will be discussed in the context of cancer cell biology and the crosstalk between Mb-dependent oxidant production and antioxidant therapeutic strategies for breast cancer.

## Material and Methods

### Cell Culture

MDA-MB-231 (CRM-HTB-26) and MDA-MB-468 (HTB-132) were purchased from The American Type Culture Collection (ATCC). MDA-MB-231 cell lines were cultured in DMEM with glutamine (4.5g/L glucose, Gibco), fetal bovine serum (FBS, 10%), 1% penicillin-streptomycin, and HEPES (pH 7.4, 15mM;). Upon initial thawing from cryopreservation, MDA-MB-231 cells were supplemented with the heme precursor δ-aminolevulinate (δ-ALA; 10 µM) for 72 hours prior to assays. MDA-MB-468 cell lines were cultured in DMEM/F-12 media (1:1; Gibco) and supplemented with FBS (10%) and penicillin-streptomycin (1%). Cells treated with mitomycin C (MMC) were treated with 20ug/mL MMC for 2 hours in complete growth media before washing out with phosphate-buffered saline (PBS, pH: 7.4) and replacing with complete growth media.

### Cell Lysis, Protein Extraction, and Estimation

Prior to lysis, cells were washed with PBS (pH: 7.4) twice before being lysed in RIPA buffer with a proteasome inhibitor cocktail and allowed to lyse for at least 20 minutes on ice. Cell suspensions were centrifuged (20,000 x g; 20 minutes; 4°C); supernatants were collected for downstream analysis. Protein concentrations were estimated using the Pierce bicinchoninic acid (BCA) Protein Assay kits following manufacturer’s instructions.

### Western Blot

Western blots were performed as previously described.^*31*^ Primary antibodies used include Mb (catalog G-125-C, R&D Systems), CPT1a (catalog 97361, Cell Signaling), and citrate synthase (catalog 14309, Cell Signaling). Blots were incubated in the appropriate fluorophore-conjugated secondary antibodies, imaged using the LI-COR Biosciences Odyssey DLx imager, and analyzed using the LI-COR Image Studio Lite (software version 5.2). All protein concentrations were normalized to the housekeeping protein α-tubulin (catalog: CP06, Calbiochem) or β-actin (catalog ab8227, Abcam) unless otherwise noted.

### Generation of Stable Myoglobin-Expressing Cell Lines via Lentiviral Transduction

MDA-MB-231 cells (90% confluence) were transduced with pLenti-C-mGFP-P2A-Puro viral particles (9.75×10^*5*^ TU; catalog: RC212352L4V; OriGene) expressing human myoglobin (*MB*; transcript variant 1; 231Mb). Empty vector control cells (231EV) were generated by transducing MDA-MB-231 cells with pLenti-C-mGFP-P2A-Puro viral particles (9.75×10^*5*^ TU; PS100093; OriGene). Cells were incubated with viral particles (18 h) and then placed under puromycin selection (1µg/mL). Cells were expanded and maintained under puromycin selection media for at least three passages before experiments.

### Transient Knockdown of Endogenous Mb via siRNA Transfection

Using the Lipofectamine RNAiMAX Transfection Reagent (catalog: 13778075; Invitrogen) according to manufacturer’s instructions, MDA-MB-468 cells were treated with either 20 nmol of ON-TARGETplus non-targeting (siNT; catalog: D-001810-10-05; Dharmacon) or Mb-targeting SMARTpool siRNA (siMb; catalog: L-012057-01-0005; Dharmacon) and incubated overnight. Transfection complexes were removed 24 hours after transfection and replaced with complete growth medium. Cells were subsequently assayed between 48 and 72 hours after transfection.

### Scratch/Wound Closure Assay

Cells were seeded and achieved 95% confluence before being treated with MMC the following day. Cells were “scratched” using a P200 pipet tip, washed with PBS, and given complete growth media. Cells in the same field of view were immediately imaged under a brightfield microscope at t=0, 2, 4, 6, or 12 hours using a reference line on the cell culture plate. The percentage of scratch/wound closure was calculated by measuring the area of the wound size via ImageJ and a ratio of the wound size at 0 hours. Data were fitted using a simple linear regression to calculate the scratch/wound closure rate.

### Transwell (Boyden Chamber) Migration Assay

Cells were serum-starved (0.5% FBS) overnight and then disassociated and added to the upper chamber of Corning Transwell polycarbonate membrane cell culture inserts (24-well format with 6.5mm inserts, 8.0-micron pore sizes). Full growth media (10% FBS) or low serum media (0.5% FBS) was added to the lower chamber of the plate as a chemoattractant or an undirected migration control, respectively, and incubated for 6 hours. In selected experiments, N-acetyl cysteine (NAC; 1mM) was added to the cell suspension in the upper chamber immediately prior to starting the assay. Cells were collected by fixing in methanol and stained with crystal violet (0.5%). Cells that did not migrate to the lower plane of the membrane were removed using a cotton-tipped applicator and allowed to dry overnight. Migrated cells were imaged under a Zeiss brightfield microscope with a 5x objective in 5 unique fields of view per transwell insert. Migrated cells were quantified using a custom ImageJ software plugin.

### Seahorse Extracellular Flux (XF) Analysis

MDA-MB-231 and MDA-MB-468 cells (25,000/well) were seeded into XF96 cell culture microplates, and oxygen consumption rate (OCR) and extracellular acidification rate (ECAR) was measured by Seahorse XFe96 Analyzer (Agilent). For mitochondrial stress tests, DMEM, oligomycin A (Oligo, 2.5µM; MilliporeSigma), FCCP (1.0µM for MDA-MB-231 cells; 0.75µM for MDA-MB-468 cells; Cayman Chemical), and rotenone (2.5µM; MilliporeSigma) were injected sequentially. For substrate-dependency assays, etomoxir (40µM for MDA-MB-231 cells; 100µM for MDA-MB-468 cells; Cayman Chemical), UK5099 (10µM; MilliporeSigma), and BPTES (10µM; Cayman Chemical) were added prior to injection of oligomycin, FCCP, and rotenone. Rates of respiration were calculated as previously described.^*32*^

### Lipidomic HPLC-MS/MS

One million cells were collected by scraping into 80% methanol and then frozen at -80C before analysis by HPLC-MS/MS as previously described.^*33*^

### Palmitate Oxidation

Complete oxidation of ^14^C-labeled palmitic acid (C_16_) was measured as previously described.^*34*^

### Transient Expression of Mb Constructs via Plasmid Transfection

MDA-MB-231 cells were transfected with Mb DNA using TransIT-X2 transfection reagent following manufacturer’s instructions (Mirus Bio) and assayed 48-72 hours after transfection.

### Generation of Apo-Mb and Mb K4M Mutants and Plasmid Expression

Apo-Mb (Mb H94F) was generated as previously described.^*15*^ To generate the Mb K46M mutant, site-directed mutagenesis (SDM) of Mb was achieved using the Phusion SDM Kit (catalog: F541; Thermo Scientific) and the parent Mb (WT) plasmid (catalog: RG212352; OriGene) according to manufacturer’s instructions and custom primers with 5’ phosphorylation modifications (Table 1). Plasmids were transformed into competent DH10B *E. coli* (catalog: C3019; New England Biolabs) and selected for ampicillin resistance according to the manufacturer’s instructions. Bacterial colonies were isolated and cultured in LB Miller Broth overnight with appropriate selection antibiotics. Plasmid DNA was isolated using QIAprep Spin Miniprep Kit (catalog: 27104; Qiagen) following the manufacturer’s instructions. Mutant Mb sequence alignments were validated using Sanger Sequencing and NCBI’s Basic Local Alignment Search Tool (BLAST). MDA-MB-231 cells were transiently transfected with Mb-expressing constructs as above and assessed 48-72 hours post-transfection. Mb protein levels were verified by western blot as above.

**Table 1:**
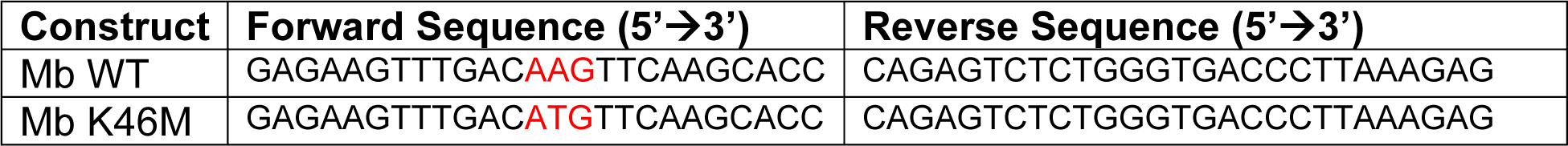
Mb K46X Mutagenesis Primers.

### Recombinant Protein Expression of Mb

Human Mb constructs (WT Mb, Mb K46M) were cloned into pET28a(+) vectors with bacterially optimized codons and purchased from GenScript Biotech Corp. pET28 plasmids were transformed into SHuffle® T7 Express competent *E. coli* (catalog: C3029J; New England Biolabs) according to the manufacturer’s instructions. Single bacterial colonies were selected, expanded, and stored as 50% LB Miller Broth/ glycerol stocks (v/v). Expression and purification of WT and mutant Mb proteins were carried out by HPLC as described previously.^*35*^

### Lipid Binding Experiments

The binding of sodium oleate to WT Mb and mutant Mb K46M constructs was assessed by UV-Vis spectroscopy as previously described.^*36*^ We verified that WT Mb and Mb K46M recombinant proteins were 80-81% and 73-77% oxy-Mb, respectively, by spectra deconvolution analysis.^*13*^ ORIGIN software (version 8) was used to fit data using *Equation 1* to determine the dissociation constant (*K_D_*) as previously described.^*36*^

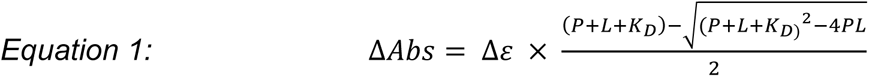

### Cellular Thermal Shift Assay (CETSA)

MDA-MB-231 cells were plated and transiently transfected with either Mb (WT), H94F (Apo-Mb), or Mb K46M plasmids as above and treated with BSA-conjugated palmitate (BSA-PA; 25µM) for 24 hours. Cells were harvested as previously described with some modifications.^*37*^ Briefly, cells were harvested and washed in PBS supplemented with protease inhibitors (PBS+PI) and then resuspended PBS+PI. Cell suspensions were freeze-thawed three times and centrifuged (20,000 x g; 20 minutes; 4°C); supernatants were collected for downstream analysis. Cell lysates were heated across a range of temperatures (40-95°C; 5 min) followed by cooling (4°C; 5 minutes) on a BIO-RAD C1000 Thermocycler. Heated samples were centrifuged (18,000 x g; 20 minutes; 4°C) to separate soluble from precipitated proteins. Supernatants were then analyzed by western blot as above. For each thermal denaturation curve, protein concentrations were normalized to 47.5°C samples and expressed as a percent of the 47.5°C signal. Melting temperature (T_m_) was determined by graphing protein signals and the corresponding temperatures as a Sigmoidal dose-response with a least-squares regression line fit using ORIGIN software, where T_m_ is equivalent to logEC_50_.

### Transient Expression of CPT1a via Plasmid Expression

Competent DH10B *E. coli* (catalog: C3019; New England Biolabs) were transformed with the Myc-DDK-tagged CPT1a transcript variant 1 (catalog: RC221740; OriGene) according to manufacturer’s instructions and colonies were selected via kanamycin resistance. Bacterial colonies were isolated and cultured in LB Miller Broth overnight with appropriate selection antibiotics. Plasmid DNA was isolated using QIAprep Spin Miniprep Kit (catalog: 27104; Qiagen) following the manufacturer’s instructions. MDA-MB-231 cells were then transiently transfected with GFP- or CPT1a-expressing constructs using the MegaTran 2.0 transfection reagent (catalog: TT210002; OriGene) according to manufacturer’s instructions. CPT1a protein levels and cell migration were assessed 48-72 hours post-transfection via western blot and Transwell assays, respectively, as above.

### Amplex™ Red H_2_O_2_ Measurement

Relative hydrogen peroxide (H_2_O_2_) production rates in MDA-MB-231 cells were measured by Amplex™ Red as previously described.^*15*^

### Pegylated Catalase (PEG-Catalase) Treatment

MDA-MB-231 cells were seeded to achieve 60% confluence, and the following day, cell media was aspirated from the plate and followed one wash with 1X PBS. Cells received full growth media with no treatment (+NT) or PEG-Catalase at a final concentration of 100U/mL. Cells were incubated for 24 hours before RNA isolation or lysis, as detailed below.

### RNA Isolation and cDNA synthesis

Cell media was aspirated from the plate and followed with three washes with 1X PBS. RNA was extracted from the cell suspension using TRIzol reagent (catalog: 15596026; Invitrogen), and nucleic acids were extracted using phenol-chloroform. Samples were centrifuged for 12,000 x g for 10 minutes to separate organic and aqueous phases. The aqueous phase was applied to RNeasy kit columns (catalog: 74104; Qiagen) and used to isolate and wash isolated RNA. Genomic DNA was eliminated, and mRNA was reverse transcribed into cDNA using the RT^*2*^ First Strand Kit according to the manufacturer’s instructions (catalog: 33040; Qiagen).

### Cell Motility qPCR Array

The Cell Motility RT^*2*^ Profiler PCR Array (catalog: PAHS-128ZE; Qiagen) was run on an Applied Biosystems Quant Studio 5 machine using RT^*2*^ SYBR Green ROX Mastermix (catalog: 330521; Qiagen) according to the manufacturer’s instructions. Gene expression was normalized to β2-microglobulin (*B2M*) for each cell line and treatment group. ΔCt values were calculated as the difference between the cell motility gene and the control gene (*B2M*), and fold change (FC) was calculated as the quotient of the 2^-ΔCt^ of the experimental groups (231Mb +NT or 231Mb +Catalase) divided by the 2^-ΔCt^ of the control group (231EV +NT). Genes in 231Mb +NT-treated cells with ≥0.20-fold change (FC) compared to 231EV +NT cells were included for further analyses if catalase treatment reverted gene expression FC by ≥0.10 towards the 231EV +NT control cell levels. Expression of *EZR*, *ITGB1*, and *RHOA* was validated with independent primers (Table 2) and samples with RT^*2*^ SYBR Green ROX Mastermix as above.

**Table 2:**
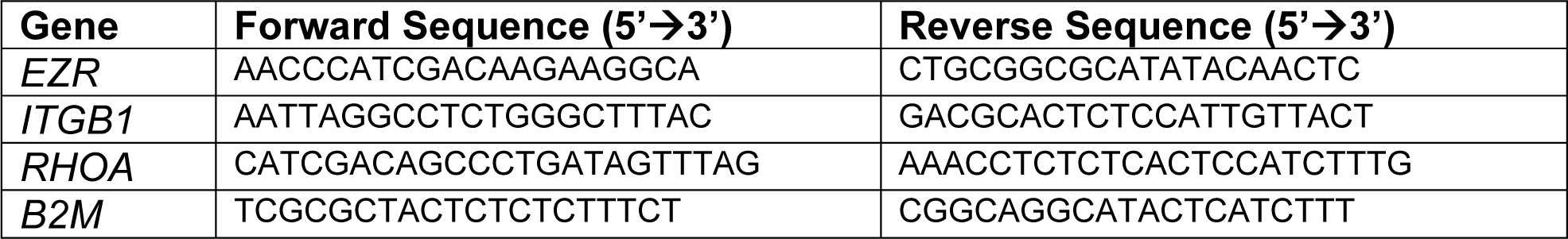
qPCR Target Validation Primers.

### Statistical Analyses

All statistical analyses were carried out using GraphPad Prism version 9.5 unless otherwise noted. Welch’s t-test was used to determine the significance between two groups for Mb-knockdown in MDA-MB-468 cells, ^14^C-palmitic acid oxidation, and lipidomics. One-way ANOVA with multiple comparisons was used to determine the significance between CETSA T_m_, Transwell, and ROS production assays with 231+GFP, 231+Mb, 231+Apo-Mb, and 231+Mb K46M cells, as well as all gene expression analyses. Two-way ANOVA with multiple comparison tests was used for Transwell migration assays, wound closure assays, Seahorse XF experiments, CPT1a protein quantification, and ROS production with NAC treatments.

## Results

### Mb expression decreases migration rates of MDA-MB-231 breast cancer cells

To determine whether Mb expression modulates breast cancer cell migration, human Mb was stably expressed in MDA-MB-231 breast cancer cells, which do not express endogenous Mb (**Figure 1A**). In a transwell migration assay, stable expression of GFP-tagged Mb (231Mb cells) significantly decreased the migration rate compared to GFP empty-vector control cells (231EV; **Figure 1B-C**). To ensure that differential proliferation was not responsible for the difference in migration, we repeated the assay in the presence of the proliferation inhibitor mitomycin C (+MMC; 20µg/mL). In the absence of proliferation, Mb expression still significantly decreased the migratory capacity of 231 cells (**Figure 1B-C).** This effect was recapitulated in a scratch closure assay in which Mb expression significantly decreased the closure of the scratch over six hours (**Figure 1D-E**). These data demonstrate that Mb expression decreases breast cancer cell migration.

**Figure 1:**
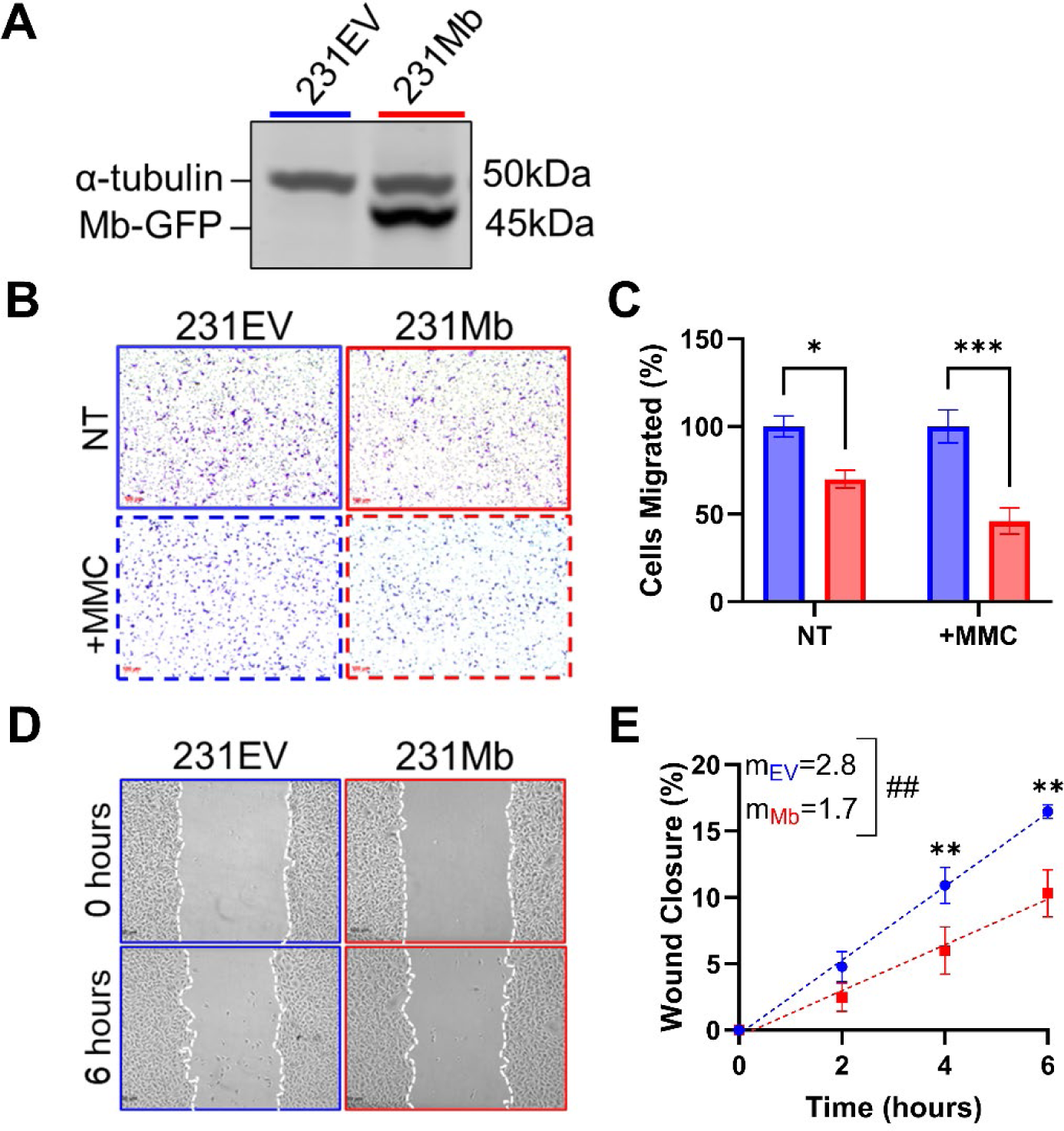
Mb expression decreases migration in MDA-MB-231 cells. **A**) Representative western blot of stable MDA-MB-231 cell lines expressing either an empty vector GFP (231EV) or GFP-tagged Mb (231Mb). **B**) Representative brightfield images of migrated cells from Transwell experiments stained with crystal violet and **C**) quantification of the percentage of 231Mb cells migrated (red bars) relative to the 231 EV cells (blue bars) at 6 hours with mitomycin C treatment (+MMC) or no treatment (NT). Two-way ANOVA with multiple comparisons tests; N=4. **D**) Representative images of 231EV and 231Mb cells in a scratch assay at 0 and 6 hours; dashed lines outline wound size. **E**) Quantification of assays such as in Panel D showing the average percentage wound closure relative to the initial wound size (t=0) and fit using a simple linear regression (dashed lines) to determine the rate of migration/wound closure (m). Simple linear regression with a comparison of slopes (**##**: p=0.0054); N=3. Two-way ANOVA with multiple comparisons test; N=3. Data are Mean ±SEM. ✱: p<0.05, ✱✱: p<0.01, ✱✱✱: p<0.001.

### Mb expression decreases the mitochondrial oxygen consumption rate (OCR) of breast cancer cells

To determine whether the Mb-induced change in migration was associated with bioenergetic alterations, we measured cellular bioenergetics in each cell line using Seahorse extracellular flux analysis. Relative to the 231EV basal oxygen consumption rate (OCR), 231Mb cells showed a significant decrease in basal (64±13%), ATP-linked (53±10%), and maximal (121±19%) OCR (**Figure 2A-B)**. We confirmed the OCR-limiting effect of Mb expression in a second breast cancer cell line, MDA-MB-468, which endogenously expresses Mb. These cells were treated with siRNA-targeting Mb, which resulted in a ∼50% decrease in endogenous Mb levels compared to non-targeting siRNA controls (**Figure 2C**). Measurement of bioenergetics in these cells showed that cells depleted of endogenous Mb (468siMb) showed a significantly increased basal and maximal OCR compared to endogenous Mb-containing cells treated with non-targeting siRNA (468siNT; **Figure 2D-E**). These data demonstrate that the expression of Mb significantly decreases the mitochondrial basal and maximal OCR in breast cancer cells.

**Figure 2:**
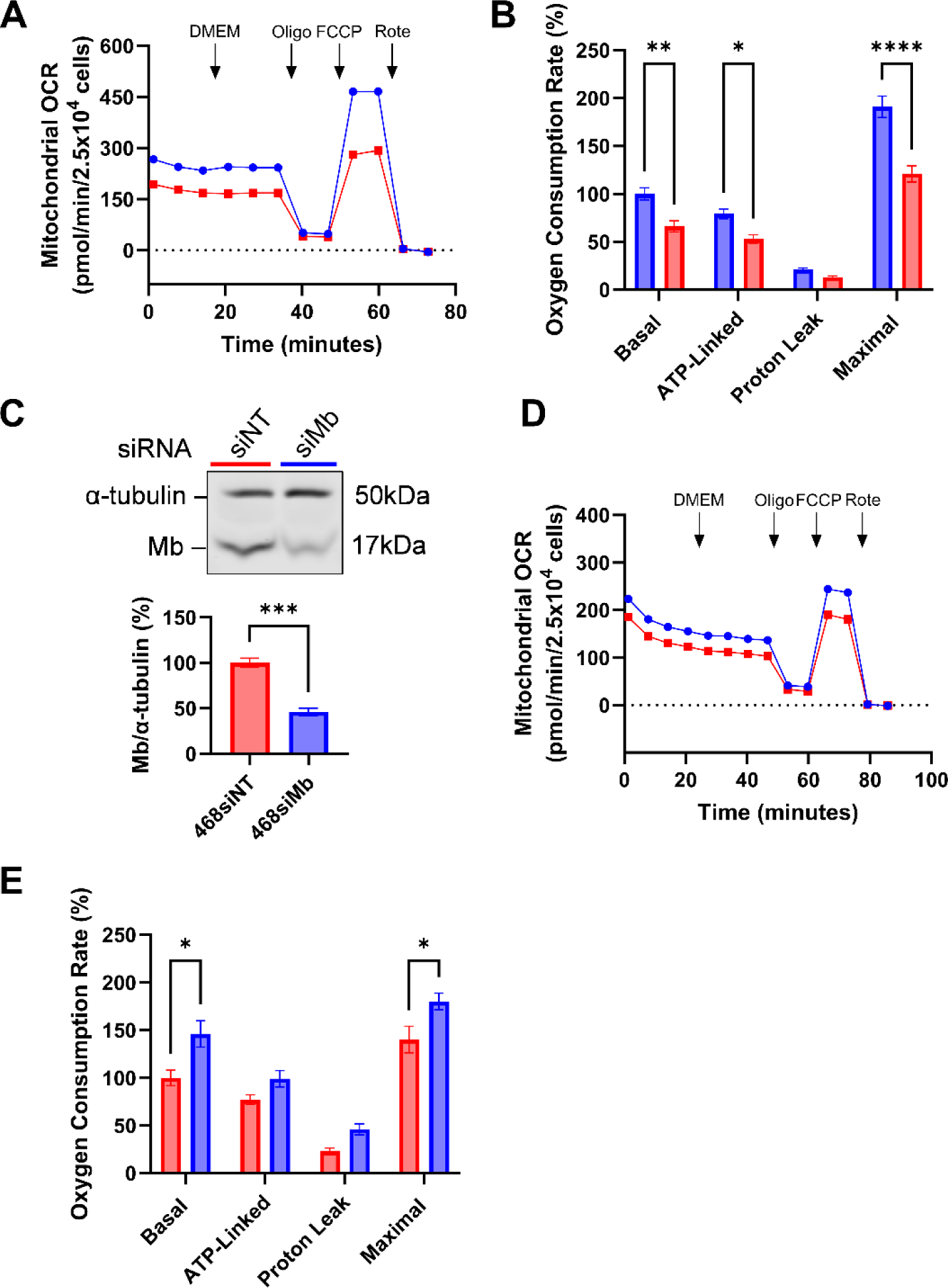
Mb expression decreases mitochondrial oxygen consumption rate (OCR). **A)** Representative Seahorse extracellular flux (XF) analyzer traces of mitochondrial OCR in 231 EV (blue dots) and 231 Mb (red squares) cells. Arrows indicate the injection of either DMEM, Oligomycin A (Oligo; 2.0µM), FCCP (1.0 µM), or Rotenone (Rote; 2.5µM). **B)** Quantitation of mitochondrial OCR in 231 EV (blue bars) and 231 Mb (red bars). Two-way ANOVA with multiple comparisons tests; N=5. **C**) Top: Representative western blot of MDA-MB-468 cells transfected with non-targeting siRNA (siNT) or Mb-targeting siRNA (siMb). Bottom: Quantitation of siRNA-mediated knockdown of Mb in MDA-MB-468 cells. Welch’s t-test; N=4. **D)** Representative traces of mitochondrial OCR in 468siNT (red squares) and 468siMb (blue dots) cells. **E)** Quantitation of mitochondrial OCR of 468siNT (red bars) and 468siMb (blue bars). Two-way ANOVA with multiple comparisons tests; N=3. Data are Mean ±SEM. ✱: p<0.05, ✱✱: p<0.01, ✱✱✱✱: p<0.0001.

### Mb expression decreases fatty acid oxidation (FAO)

To determine whether alterations in substrate utilization contribute to the Mb-dependent decrease in basal and maximal OCR, we next assessed the contribution of pyruvate, fatty acid, and glutamine oxidation to basal respiration by measuring OCR in the presence of inhibitors of the mitochondrial pyruvate carrier (UK5099: 20µM), mitochondrial fatty acid entry through carnitine palmitoyl transferase-1 (CPT1; etomoxir: 40µM), or glutaminase (BPTES: 20µM; **Figure 3A-B**). The contribution of pyruvate and glutamine utilization to OCR was not statistically different in 231Mb cells compared to 231EV cells (**Figure 3B**). However, inhibition of mitochondrial fatty acid import using etomoxir showed a significant decrease in fatty acid-dependent OCR in 231Mb cells (28±6%) compared to 231EV cells (41±6%; **Figure 3A-B**). Consistent with Mb-dependent suppression of fatty acid oxidation, 468siMb cells also showed a higher contribution of fatty acid oxidation to basal OCR than 468siNT cells (468 siNT: 86±15%, 468 siMb: 53±4%; **Figure S1**).

**Figure 3:**
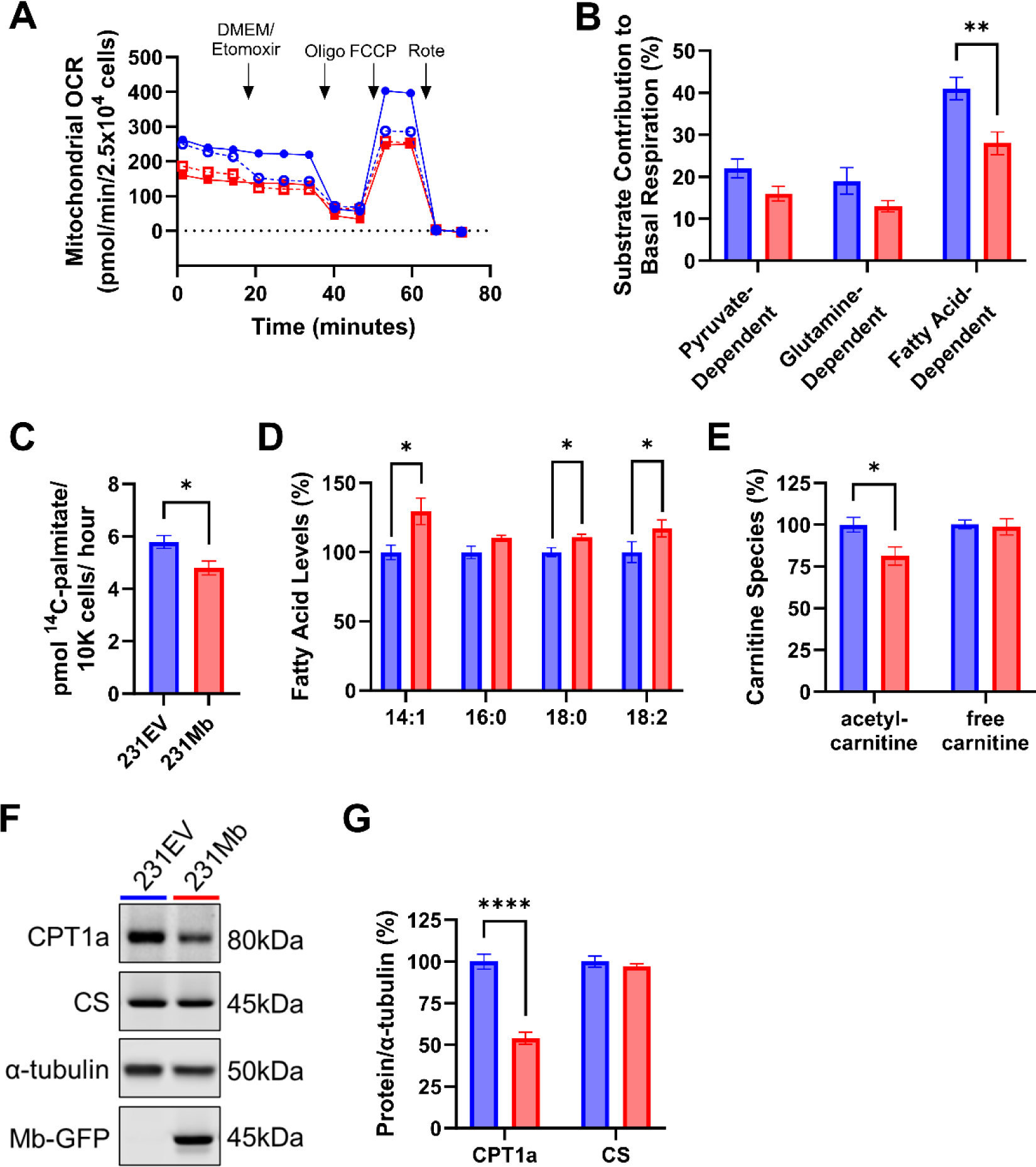
Mb expression decreases fatty acid oxidation (FAO). **A)** Representative traces of mitochondrial OCR of 231 EV (blue dots) and 231 Mb (red squares) cells. Dashed lines with open symbols indicate etomoxir-treated cells (40µM). Arrows indicate the time of injection of DMEM or Etomoxir, Oligo, FCCP, or Rote. **B)** The percent contribution of pyruvate-, glutamine-, and fatty acid-dependent respiration to total basal OCR in 231EV (blue bars) and 231Mb (red bars) cells. Two-way ANOVA with multiple comparisons tests; N=5. **C)** Rate of ^14^C-palmitic acid oxidation in 231EV (blue bars) and 231Mb (red bars) cells. Welch’s t-test; N=9-10. **D)** Levels of myristoleic (14:1), palmitic (16:0), stearic (18:0), and linoleic (18:2) fatty acids in 231 EV (blue bars) and 231 Mb (red bars) cells. Welch’s t-test; N=6. **E)** Relative levels of carnitine species extracted from 231EV (blue bars) and 231Mb (red bars) cells as measured by LC-MS. Two-way ANOVA with multiple comparisons tests; N=6. **F-G)** Representative western blots (F) and quantitation (G) of carnitine palmitoyl transferase-1a (CPT1a) and citrate synthase (CS) in 231 EV (blue) or 231 Mb (red) cells. Two-way ANOVA with multiple comparisons tests; N=3-4. Data are Mean ±SEM. ✱: p<0.05, ✱✱: p<0.01, ✱✱✱✱: p<0.0001.

To confirm these results, we directly assayed fatty acid oxidation (FAO) by measuring the oxidation of ^14^C-palmitate (C_16_) in 231EV and 231Mb cells. Mb expression significantly decreased palmitate oxidation in the 231Mb cells (4.797±0.838 pmol/hour) compared to the 231EV cells (5.789±0.729 pmol/hour; **Figure 3C**). Furthermore, using liquid chromatography-coupled mass spectrometry (LC-MS), we observed that levels of myristoleic (C14:1), palmitic (C16:1), stearic (C18:0), and linoleic (C18:2) fatty acids were elevated (between 11-29%) in 231Mb cells relative to 231EV cells (**Figure 3D**). Levels of free carnitine were not changed between cell lines, but acetyl-carnitine was significantly decreased in 231Mb cells, consistent with decreased FAO in 231Mb cells (**Figure 3E**). We next examined the levels of CPT1a, which catalyzes the rate-limiting step of FAO.^*38*^ Consistent with Mb decreasing FAO, 231Mb cells show a 46±7% decrease in CPT1a levels compared to 231EV cells **(Figure 3F-G**). We show that citrate synthase (CS), a marker of mitochondrial mass,^*39*^ is not changed with Mb expression **(Figure 3F-G**), suggesting that Mb expression does not affect mitochondrial size. Collectively, these data show that Mb expression significantly decreases FAO, leading to increased free fatty acid levels in MDA-MB-231 cells.

### Inhibition of Mb-dependent fatty acid binding and overexpression of CPT1a do not restore migration rates of Mb-expressing cells

Computational models of Mb-fatty acid binding suggest key electrostatic interactions between the positively charged lysine residue (K45) of equine oxy-Mb and the negatively charged carboxylate head of palmitic and oleic acids, while the alkyl tail of the fatty acid tucks into a hydrophobic cleft near the heme prosthetic group of oxyMb.^*24*, *25*, *40*^ To test whether Mb-dependent attenuation of FAO is due to its ability to bind fatty acids, we generated a mutant Mb protein in which the K46 (the human ortholog of equine Mb K45) was mutated from lysine to methionine (K46M). UV-Vis spectroscopy of these recombinant proteins demonstrated no change in the Soret peak (∼410nm) of the Mb K46M mutant construct compared to wildtype (WT) Mb, demonstrating that heme incorporation is unaffected by mutagenesis (**Figure 4A**). Increasing concentrations of oleate (18:1; 0-200µM) decreased the absorbance of the Soret peak of both the WT and MbK46M proteins, consistent with oleate binding (**Figure 4B**). Calculating the differential spectra and fitting the kinetic binding curves showed distinctly different fatty acid binding profiles for WT and K46M Mb (**Figure 4C-D**). Calculation of the equilibrium constant (K_D_) showed a ∼50-fold increase in the K_D_ of Mb K46M (13.13±6.75µM oleate) compared to that of WT Mb (0.24±0.10µM oleate), suggesting that the Mb K46M mutant does not stably bind oleic acid. These results were confirmed by a secondary assay of ligand binding, Cellular Thermal Shift Assay (CETSA), in which binding of fatty acid to Mb was expected to stabilize the protein and increase its melting temperature (T_m_). Consistent with the disruption of Mb-fatty acid binding, the T_m_ for Mb was significantly decreased in cells expressing Mb K46M compared to WT Mb (**Figure 4E & S2**).

**Figure 4:**
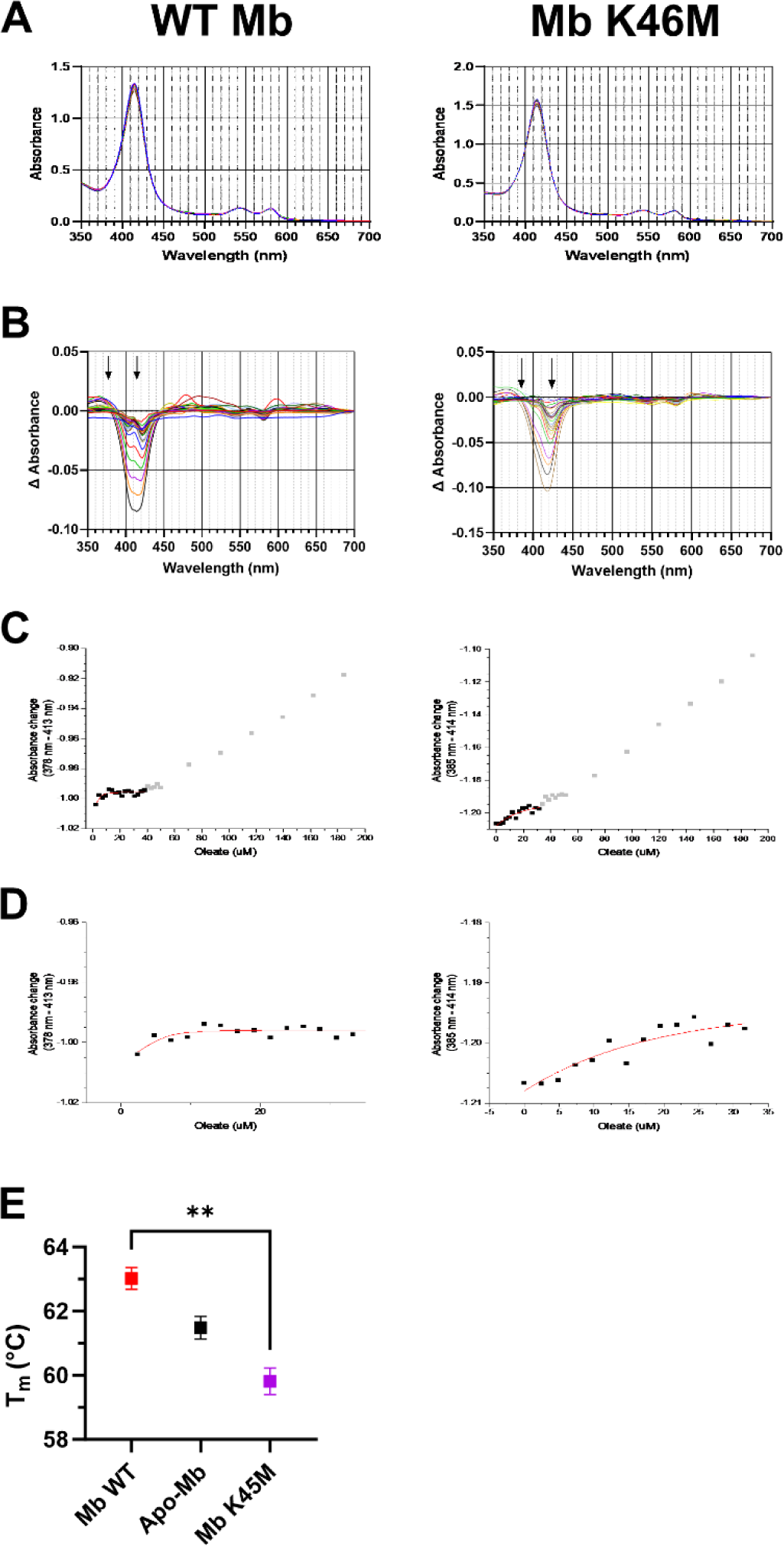
Mutation of Mb K46 to methionine (K46M) is sufficient to decrease Mb’s affinity for oleate. **A**) Representative absorbance spectra of WT Mb (left) and Mb K46M (right). **B**) Representative differential spectra of WT and K64M Mb with increasing concentrations of oleate (0-200µM). Arrows represent isosbestic and Soret peak wavelengths used to generate binding curves. **C-D**) Representative absorbance changes for each protein as a function of oleate concentration and data fit. C is the representative data fit using the low oleate concentrations (black squares) using a binding model of 1:1 stoichiometry based on the equilibrium constant and Beer-Lambert equation (Equation 1). Measurements at high oleate concentrations (grey squares) were not used in the data fit and calculation of the binding constant. N=2. **E**) Calculated T^m^ of WT Mb (red), Apo-Mb (black), or Mb K46M (purple) protein from CETSA in MDA-MB-231 cells. One-way ANOVA with multiple comparisons to the T_m_ of WT Mb; N=3. Data are Mean ±SEM; ✱✱: p=0.0028.

To test whether elimination of Mb-fatty acid binding attenuated the Mb-dependent decrease in mitochondrial OCR, we transfected 231 WT cells with either a GFP-control, WT Mb, or Mb K46M plasmids (231+GFP, 231+Mb, 231+Mb K46M, respectively) and measured bioenergetics (**Figure 5A-B**). As before, 231+Mb cells showed decreased basal, ATP-linked, and maximal mitochondrial OCR compared to control cells (231+GFP, which express no Mb). This effect was potentiated, rather than attenuated, in the 231+Mb K46M mutant cells (**Figure 5A-B**). Further, compared to the 231+GFP cells, 231+Mb K46M cells showed a similar decrease in migration (67±10%) as the 231+Mb expressing cells (65±9%; **Figure 5C-D**). Taken together, these data demonstrate that Mb decreases OCR and migration through a mechanism independent of Mb’s ability to bind fatty acids.

**Figure 5:**
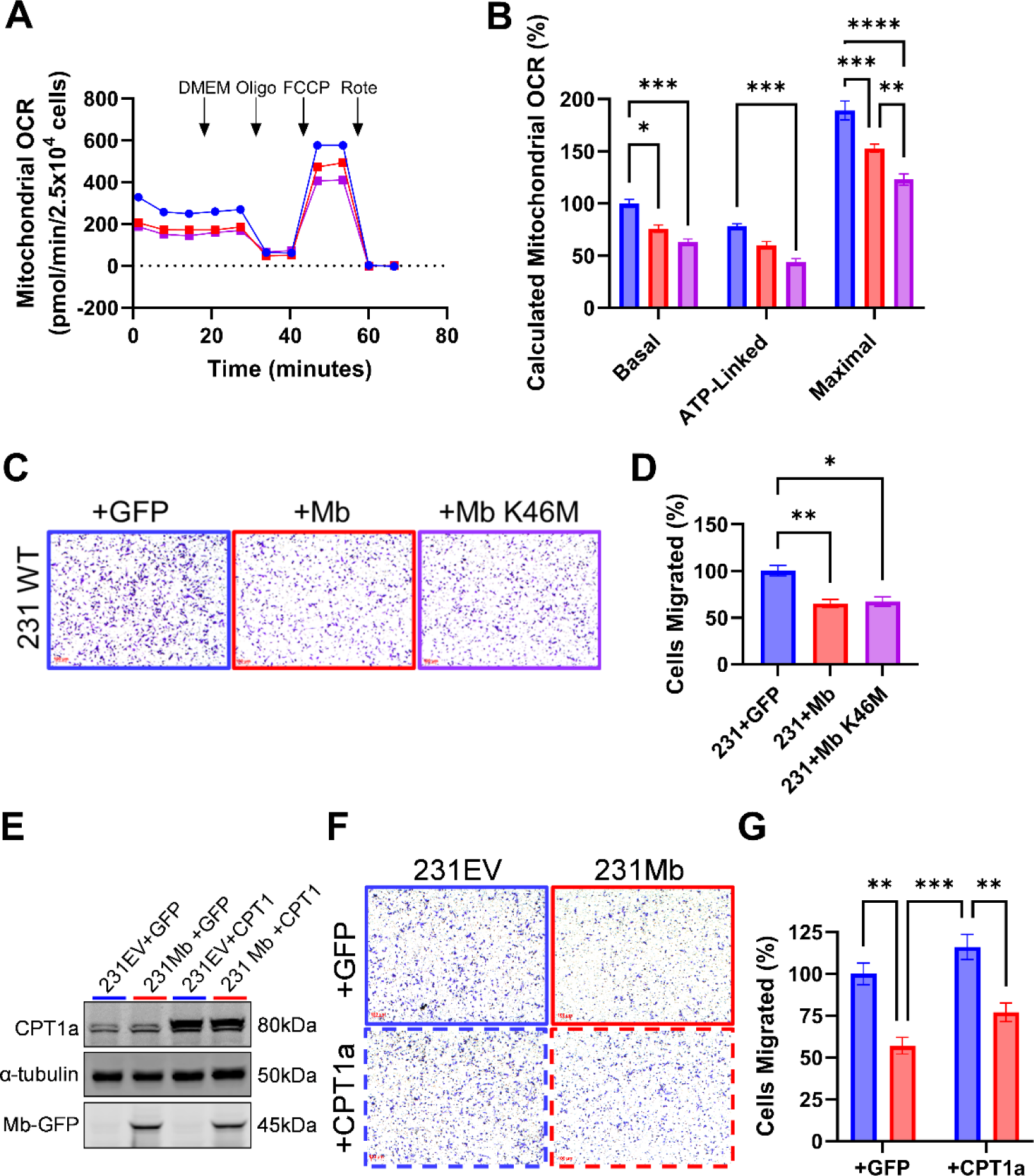
Elimination of fatty acid binding and overexpression of CPT1a is not sufficient to attenuate Mb-dependent decreases in mitochondrial OCR or migration. **A-B)** Representative mitochondrial OCR traces of cells expressing GFP vector (blue dots), WT Mb (red squares), or Mb K46M (purple squares) (A) and corresponding quantitation (B) of MDA-MB-231 WT cells transiently expressing either GFP (blue bars), WT Mb (red bars), or Mb K46M (purple bars). Arrows in Panel A indicate the time of injection of either DMEM, Oligo, FCCP, or Rote. Two-way ANOVA with multiple comparisons tests; N=4. **C)** Representative brightfield images of migrated cells from 6-hour Transwell experiments stained with crystal violet. **D)** Percent of GFP-(blue), WT Mb-(red), or Mb K46M-(purple) expressing cell migration at 6 hours. One-way ANOVA with multiple comparisons tests; N=4. **E)** Representative western blots of CPT1a levels 72 hours after transfection with control-(+GFP, solid frame) or CPT1a (+CPT1a, dashed frame) -encoding plasmids in 231EV (blue) or 231Mb (red) cells. **F)** Representative brightfield images of migrated cells from 6-hour Transwell experiments stained with crystal violet. **G)** Percent of 231EV (blue bars) and 231Mb (red bars) cells transiently expressing GFP or CPT1a at 6 hours. Two-way ANOVA with multiple comparisons tests; N=4. Data are Mean ±SEM; ✱: p<0.05, ✱✱: p<0.01, ✱✱✱: p<0.001, ✱✱✱✱: p<0.0001.

To determine whether the Mb-dependent decrease in migration was linked to the observed decrease in CPT1 expression, we overexpressed CPT1a (**Figure 5E**) and assayed migration. Overexpression of CPT1a did not significantly change the migration rate of 231Mb cells (**Figure 5F-G**), suggesting that the Mb-dependent inhibition of migration is likely independent of changes in FAO.

### Mb-dependent oxidant production decreases mitochondrial respiration and cell migration

Given that fatty acid binding was not responsible for the Mb-dependent decrease in bioenergetics and cell migration, we next tested whether the heme moiety played a role in this effect. We previously generated a mutant Mb protein in which the distal histidine (H94) was replaced with phenylalanine (H94F; heretofore referred to as “Apo-Mb”);^*15*^ Apo-Mb is unable to coordinate heme. 231 WT cells transiently expressing Apo-Mb (231+ApoMb) showed a restoration of basal, ATP-linked, and maximal OCR compared to 231+Mb cells (**Figure 6A**). Similarly, in a wound closure assay, 231+Mb cells showed significantly decreased scratch closure compared to 231+GFP cells. However, 231+ApoMb cells showed no significant difference in scratch closure compared to 231+GFP cells (**Figure 6B-C**). These data demonstrate that Mb-dependent attenuation of migration is dependent on the Mb heme moiety.

**Figure 6:**
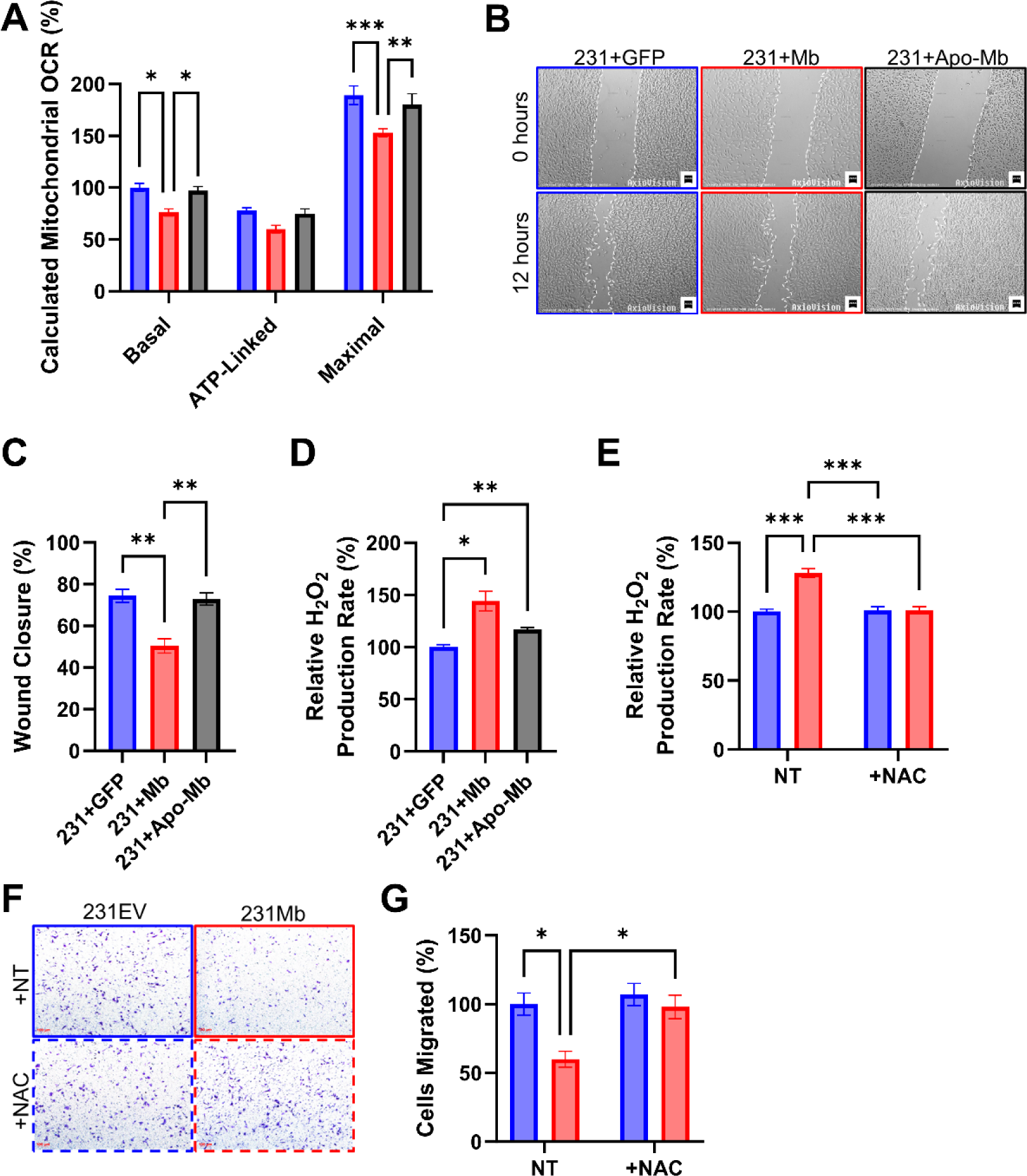
Mb-dependent oxidant production downregulates mitochondrial OCR and migration. **A**) Quantitation of OCR of MDA-MB-231 WT cells transiently expressing either GFP (blue bars), WT Mb (red bars), or Apo-Mb (black bars). One-way ANOVA with multiple comparisons tests; N=4. **B-C**) Representative brightfield images of scratch assays of 231WT with either GFP (blue frame), Mb (red frame), or Apo-Mb (black frame), and results (**C**) were quantified as a percentage relative to the initial wound size (t=0) after 12 hours with no treatment (+NT) or with catalase (+Catalase). Two-way ANOVA with multiple comparison tests; N=4 **D**) Cellular H_2_O_2_ production rates of 231WT cells expressing either GFP (blue bars), WT Mb (red bars), or Apo-Mb (black bars). One-way ANOVA with multiple comparisons tests; N=3. **E**) Cellular H_2_O_2_ production rates of 231EV (blue bars) and 231Mb (red bars) cells with no treatment (NT) or treated with N-acetyl cysteine (1mM, +NAC). Two-way ANOVA with multiple comparisons tests; N=3. **F**) Representative brightfield images at 6 hours of migrated 231EV (blue frame) and 231Mb (red frame) that are either not treated (NT, solid frame) or treated with NAC (dashed frame). **G**) Quantitation of Transwell migration assay of 231EV (blue) and 231Mb (red) cells with or without NAC treatment at 6 hours. Two-way ANOVA with multiple comparisons tests; N=3. Data are Mean ±SEM. ✱: p<0.05, ✱✱: p<0.01, ✱✱✱: p<0.001.

It is well established that in addition to binding oxygen, the heme center of Mb can catalyze oxidant production.^*15*, *41*, *42*^ Consistent with heme-dependent oxidant production, 231+Mb cells generated significantly more H_2_O_2_ than 231+GFP cells (**Figure 6D**). This effect was attenuated in 231+Apo-Mb, which lacks heme coordination (**Figure 6D**). To determine whether Mb-dependent oxidant production led to decreased cell migration, 231EV and 231Mb cells were treated with the antioxidants pegylated-catalase (PEG-catalase; +Catalase) or N-acetylcysteine (+NAC). Treatment with catalase increased the migration rates of the 231+Mb cells exclusively, and no significant changes in migration were observed in 231+GFP or 231+Apo-Mb cells (**Figure 6C**). Treatment with NAC decreased H_2_O_2_ production to a level indistinguishable from 231EV cells (**Figure 6E**). Similar to scratch assay results, Transwell migration assays demonstrated that while 231Mb cells showed significantly decreased migration, NAC treatment reversed this effect (**Figure 6F-G**). Collectively, these data suggest that Mb-derived oxidants result in decreased cellular migration.

### Mb-catalyzed oxidant generation dysregulates the transcription of key focal adhesion genes

We next sought to determine whether Mb-dependent oxidants decrease cancer cell migration by targeting proteins that regulate actin skeleton rearrangement. We measured the mRNA levels of key migratory proteins in 231EV and 231Mb cells in the presence or absence of PEG-catalase to scavenge H_2_O_2_ (**Figure 7A**). Cell motility genes that showed a fold change (FC) ≥0.20 in 231Mb non-treated (231Mb +NT) compared to 231EV +NT were considered only if PEG-catalase (+Catalase) treatment reverted gene expression changes by FC ≥0.10. Our analysis revealed 11 genes (*ACTR2*, *EZR*, *FGF2*, *HFG*, *IGF1*, *ILK*, *ITGA4*, *ITGB1*, *RHOA*, *TLN1*, and *WASF1*) that are altered in 231Mb cells and trend towards returning to the levels of 231EV +NT cell levels after catalase treatment (**Figure S3**). Of these 11 genes, *EZR*, *ITGB1*, and *RHOA* were decreased in 231Mb cells and restored with catalase (**Figure 7B**) in validation studies with independent primers (Table 2). Changes in these genes are significant as *EZR* and *ITGB1* each contribute to cell migration by linking the actin cytoskeleton to the plasma membrane^*43*–*45*^ and extracellular matrix,^*46*, *47*^ respectively. *RHOA* promotes actin polymerization and reorganization at lamellipodia to initiate membrane ruffling, an early step in cell migration.^*48*, *49*^ Taken together, these data demonstrate that Mb-dependent oxidant production decreases transcription of key cell motility genes that regulate focal adhesion structure and signaling,^*50*^ which is reversed in the presence of the antioxidant catalase.

**Figure 7:**
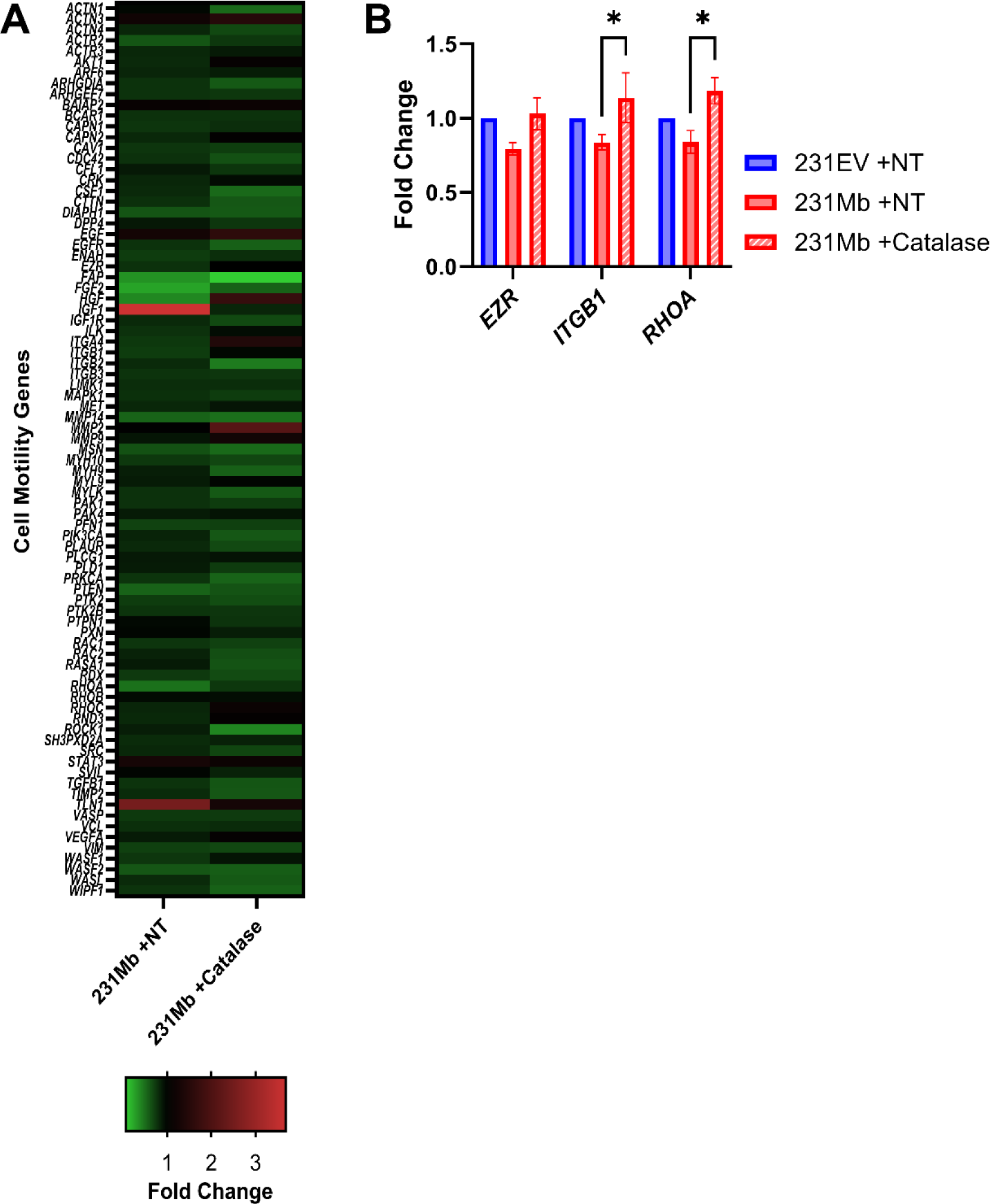
Mb expression alters the transcriptional program of migratory genes through oxidation of SP1. **A)** Heatmap gene expression fold changes from the QIAGEN cell motility RT^*2*^ Profiler array. Gene expression is represented as the fold change relative to 231EV +NT cells. Genes that are decreased are green, and genes that are increased are red. N=3-4. **B)** Independently validated gene targets *EZR*, *ITGB1,* and *RHOA* expression levels in 231Mb +NT cells (red, solid basrs) that show a ≥0.20 fold-change (FC) from the control cells (231EV +NT, blue bars) and revert to the control cell levels by FC≥0.10 with catalase treatment (red, striped bars). *EZR*: ezrin; *ITGB1*: integrin subunit beta-1; *RHOA*: ras homolog family member A. One-way ANOVA with multiple comparisons; N=3-4. ✱: p<0.05.

## Discussion

The data presented in this report demonstrate that while Mb binds to fatty acids in breast cancer cells, this interaction does not modulate mitochondrial FAO or cell migration. Instead, the Mb-dependent inhibition of respiration and cell migration is secondary to the heme-mediated generation of oxidants that modulate the expression of migratory genes. Results from our study highlight multiple cellular alterations that occur with Mb expression in MDA-MB-231 cells but center on the heme prosthetic group, rather than non-heme fatty acid binding, as the major regulator of breast cancer cell function.

Few studies have directly interrogated Mb-dependent fatty acid binding and tested whether this interaction modulates FAO in cancer cells. Christen *et al.* reported in immortalized brown adipocytes that the double-lysine mutant Mb K45A/K63A decreases Mb binding to palmitic and oleic acid. Armbruster *et al.* reported that WT Mb expression promotes cytoplasmic solubility and limits oxidation in MDA-MB-468 and MCF7 breast cancer cell lines using lipidomic measurements^*5*^, but did not directly test whether Mb-fatty acid binding is responsible for this effect. The results presented herein confirm these prior studies showing that the lysine mutation disrupts fatty acid binding and that Mb expression modulates FAO in cancer cells. However, our data demonstrate that the Mb-fatty acid binding is not responsible for Mb-dependent FAO modulation. This is likely due to the low concentration of Mb expressed in cancer cells relative to the free fatty acid concentration. Mb-fatty acid binding occurs in a 1:1 ratio^*21*–*23*^ and the concentration of free fatty acids in mammary epithelial cells is approximately five micrograms (5µg) per million cells,^*51*^ while endogenous Mb expression is between 24 and 65 nanograms (ng) per million cells.^*8*, *11*^ Therefore, the physiological levels of Mb in breast cancer cells are approximately 75-to 100-fold lower than the levels of fatty acids, suggesting that Mb expression will have negligible impact on the large pool of fatty acids available for oxidation in the cell. Surprisingly, our data suggest that in the absence of Mb-fatty acid binding, the Mb-dependent decrease in OCR is further decreased compared to wildtype Mb (rather than attenuated; **Figure 5A-B**). While this data definitively excludes a role for Mb acting as a lipid sink that binds fatty acids to prevent them from being oxidized, it is unclear whether fatty acid binding impacts Mb-dependent oxidant production, which could regulate FAO. Further study is required to determine whether the K46 mutation increases Mb-dependent oxidant production. Alternatively, fatty acids bound to Mb may serve as a target for oxidants produced at the heme site, which would scavenge these oxidants and decrease their bioavailability to modulate FAO at other sites in the mitochondrion.

Our data demonstrate that Mb expression decreases cellular respiration, consistent with prior results from Kristiansen *et al.,* who show that Mb expression decreases mitochondrial respiration in MDA-MB-468 cells.^*11*^ Our data demonstrate this decreased respiration is due to an inhibition of FAO, concomitant with decreased expression of CPT1a. CPT1 catalyzes the rate-limiting step in the FAO pathway by linking carnitine to acyl-CoAs for transport across the outer mitochondrial membrane.^*38*^ CPT1 activity has been shown to be concentration-dependently decreased by H_2_O_2_ in a manner that is reversed by catalase.^*52*^ Additionally, peroxisome proliferator-activated receptors (PPARs) drive the expression of FAO genes, including CPT1, and are downregulated by H_2_O_2_.^*53*^ These studies provide potential mechanistic pathways by which Mb heme-mediated oxidant production may downregulate CPT1 in our cell model. While our data suggest that the downregulation of CPT1 does not mechanistically underlie the decrease in cellular migration observed with Mb expression (**Figure 5F-G**), it is possible that Mb-dependent FAO downregulation attenuates other tumorigenic pathways. For example, inhibition of CPT1 has been shown to sensitize cancer cells to apoptosis, decrease ATP production for tumor cell growth, and regulate gene expression.^*54*–*58*^ In this regard, Mb expression may mimic the anti-tumorigenic effects of FAO inhibitors, which have been proposed as potential therapeutics for breast cancer.^*27*, *28*, *59*–*63*^ Future studies will test the therapeutic efficacy of FAO inhibitors on Mb-expressing tumors as well as develop strategies to enhance Mb expression in non-expressing tumors.

Our data demonstrating that Mb expression decreases MDA-MB-231 cell migration are consistent with previous studies. Aboouf *et al.* demonstrate that the knockdown of endogenous *MB* in MCF7 cells increases cell migration. However, *MB*-depleted SKBR3 and MDA-MB-468 cells showed significantly slower migration rates.^*11*, *17*^ Discrepancies in Mb’s effect on cell migration may be due to differing levels of Mb expression,^*8*, *10*^ other oxidant-generating proteins (e.g., NAPDH oxidases, lipoxygenases), and/or the levels of the heme iron-reducing proteins to allow for subsequent catalysis, like cytochrome B5 reductase 3 (CYB5R3),^*64*^ between cell lines. The expression status of hormone receptors (e.g., estrogen and progesterone) may also alter Mb’s effects on migration, as treating breast cancer cells with estradiol decreases *MB* expression.^*10*^ Further comparison between cell line models with and without Mb is required to elucidate which factors impact Mb-dependent modulation of migration.

We and others have demonstrated that Mb increases oxidant production in breast cancer cells,^*15*, *17*^ leading to dysregulated redox homeostasis and cell function.^*11*^ We have previously shown that the E3 ubiquitin ligase Parkin is oxidized in the presence of Mb, leading to changes in cell proliferation.^*15*^ Aboouf *et al*. report that Mb expression elevates oxidants and annexin-V positivity in MCF7 cells and that Mb correlates with cleaved caspase-3 in invasive breast tissues, signifying that Mb-dependent ROS contributes to apoptosis.^*17*^ Data presented here for the first time demonstrate that Mb decreases the expression of the migratory genes *EZR*, *ITGB1*, and *RHOA*. These genes encode ezrin, β1-integrin, and Ras homolog family member A (RhoA), respectively, and their protein functions converge at focal adhesions.^*50*^ Importantly, each of these genes has been shown to be elevated in various cancers and increase cancer cell migration and metastasis, contributing to worsened patient outcomes.^*43*, *45*, *65*–*75*^ Specifically in MDA-MB-231 cells, others have shown that knockdown of ezrin and β1-integrin decreases cell migration^*65*, *69*^, and knockdown of RhoA decreases cell invasion.^*75*^ Our data are in agreement with these reports in that 231Mb cells with decreased migration rates also have decreased *EZR*, *ITGB1*, and *RHOA* expression. Notably, the expression of these genes is restored with catalase treatment, demonstrating redox-sensitive regulation of gene expression. To this end, the expression of *EZR*, *ITGB1*, and *RHOA,*^*76*–*79*^ as well as particular FAO genes (including CPT1),^*80*, *81*^ is known to be regulated by the transcription factor Specificity Protein 1 (Sp1). While we did not specifically examine Sp1 in our studies, oxidants have been shown to eliminate Sp1 transcription factor activity^*82*–*84*^ and indirectly regulate Sp1 levels.^*85*^ Thus, it is interesting to speculate that Mb-dependent oxidant production decreases Sp1 activity and/or levels, leading to decreased expression of CPT1 and migration genes. Indeed, others have reported Mb alters gene expression, ^*7*, *17*^ but have not speculated on the mechanism.

It is well established that dysregulated redox homeostasis and oxidant signaling within tumors play a central role in tumor initiation, metabolic reprogramming, proliferation, metastasis, and cell death.^*86*–*89*^ Pre-clinical studies and clinical trials testing antioxidants, including vitamin C, carotenoids, and NAC, in lieu of and in addition to standard chemotherapeutics have provided mixed results.^*90*^ In breast cancer specifically, some trials have demonstrated that antioxidants limit carcinogenic oxidant signaling,^*90*, *91*^ while others show that antioxidants accelerate tumorigenesis.^*92*, *93*^ The results of our study suggest that the low levels of oxidants generated by Mb-expressing cells are central in limiting breast cancer cell FAO and migration. Thus, it is critical to understand the intratumoral sources and levels of oxidants and their regulation of cell metabolism and migration to develop effective, personalized therapies to treat metastatic breast cancers. For example, Jung and colleagues report that disease progression was increased in post-menopausal breast patients taking antioxidants (e.g., dietary vitamin C supplements) during chemotherapeutic and/or radiation therapy.^*94*^ Notably, the level of Mb expression is reported to be higher in post-menopausal breast cancer patients.^*10*^ Consequently, it is interesting to speculate that the use of antioxidants in post-menopausal women may attenuate Mb-dependent oxidant production and its downstream decrease in cell migration, driving disease progression. Future studies that stratify breast cancer patients based on Mb levels may reveal whether Mb underlies the differing effects of antioxidants on metastatic progression in breast cancer.

In conclusion, we show that Mb expression in breast cancer cells decreases mitochondrial respiration, and this is due to the inhibition of FAO and downregulation of CTP1a. This effect on respiration is concomitant with Mb-dependent attenuation of cell migration. At a mechanistic level, while Mb binds fatty acids in cancer cells, disruption of this binding does not reverse the Mb-dependent inhibition of respiration or migration. Instead, we demonstrate that the Mb heme prosthetic group increases cellular oxidant production, and these oxidants downregulate genes that regulate cellular migration. These studies define an additional role for Mb’s heme moiety in the regulation of fatty acid metabolism and migration. Further, our data underscore the role of antioxidant-oxidant signaling in cancer cell processes and highlight the importance of considering Mb expression in personalized breast cancer therapies.

## Acknowledgements

We would like to thank Dr. Courtney Sparacino-Watkins for her insight and assistance with measurements of Mb-fatty acid binding.

This work was supported by National Institutes of Health Grants R01HL166985-01A1 and support from the Hemophilia Center of Western Pennsylvania (SS) and T32 HL110849 (to ARJ).

