## Supplemental Figures for "Myoglobin Inhibits Breast Cancer Cell Fatty Acid Oxidation and Migration via Heme-dependent Oxidant Production and Not Fatty Acid Binding"

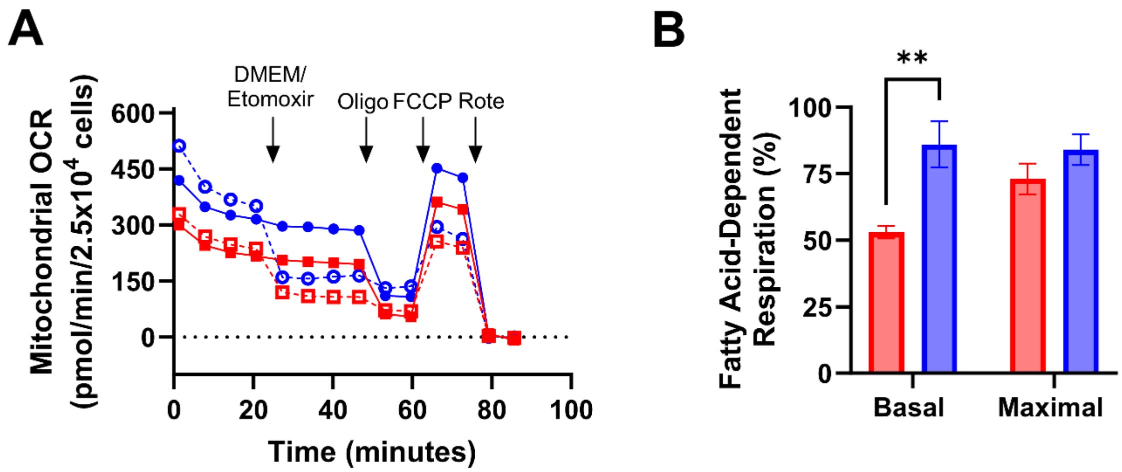


**Figure S1: Expression of Mb in MDA-MB-468 cells decreases FAO. A)** Representative traces OCR in 468siNT (red squares) and 468siMb (blue dots) cells. Dashed lines and open symbols indicate etomoxir-treated cells. Arrows indicate the time of injection of either Etomoxir, Oligo, FCCP, or Rote. **B)** The percent contribution of fatty acid-dependent respiration to total basal and maximal OCR in 468siNT (red bars) and 468siMb (blue bars) cells. Two-way ANOVA with multiple comparisons tests; N=3. Data are mean ±SEM. ✱✱: $p<0.01$.

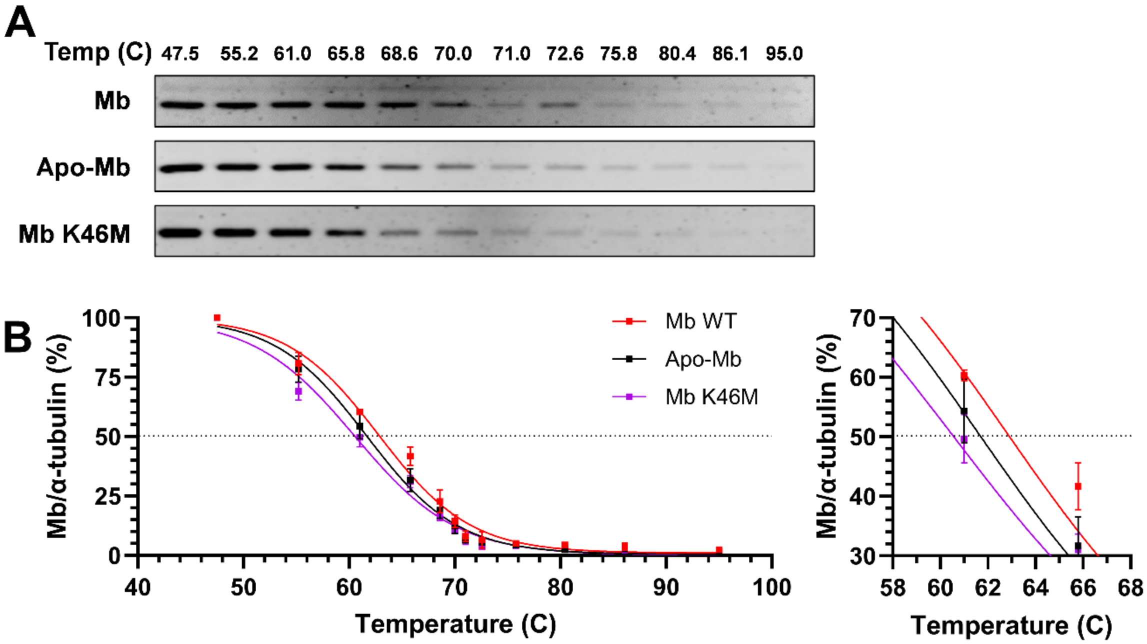


**Figure S2: Mb K46M decreases protein stability by cellular thermal shift assay (CETSA). A)** Representative western blots of MDA-MB-231 WT cells expressing either WT Mb, Apo-Mb, or Mb K46M in which protein concentration decreases as a function of increasing temperature. **B)** WT Mb (red squares), Apo-Mb (black squares), and Mb K46M (purple squares) protein levels transiently expressed in MDA-MB-231 cells decrease as a function of temperature. Protein signals are fit to a sigmoidal dose-response curve with a variable slope by least-squares at all temperatures (left) and near the melting temperature ($T_m$, dashed line; right). For all panels, N=3. Data are Mean ±SEM.

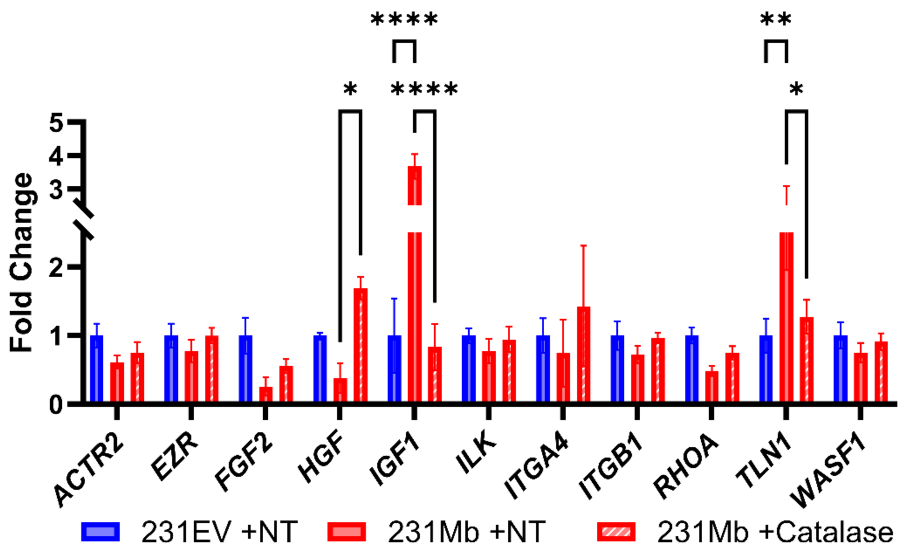


**Figure S3: Mb-dependent oxidant production causes differential gene expression.** Eleven differentially expressed genes (DEGs) implicated in cell motility are altered (FC≥0.20) with Mb expression (231Mb+NT, red, solid bars) relative to control cells (231EV+NT, blue, solid bars). Expression of these genes in 231Mb cells is increased (FC≥0.10) with catalase treatment (231Mb +Catalase, red, striped bars) relative to the 231Mb +NT cells. *ACTR2*: actin related protein 2 ; *EZR*: ezrin ; *FGF2*: fibroblast growth factor-2; *HGF*: hepatocyte growth factor; *IGF1*: insulin growth factor-1; *ILK*: integrin-linked kinase; *ITGA4*: integrin subunit alpha-4 ; *ITGB1*: integrin subunit beta-1; *RHOA*: ras homolog family member A; *TLN1*: talin-1; *WASF1*: Wiskott-Aldrich syndrome protein (WASP) family member 1. Two-way ANOVA with multiple comparisons; N=3-4. Data are mean ±SEM. ∗: $p<0.05$, ∗∗: $p<0.01$, ∗∗∗∗: $p<0.0001$.
